## Supplemental Figures for "Circadian rhythms of macrophages are altered by the acidic pH of the tumor microenvironment"

Supplementary Fig 1

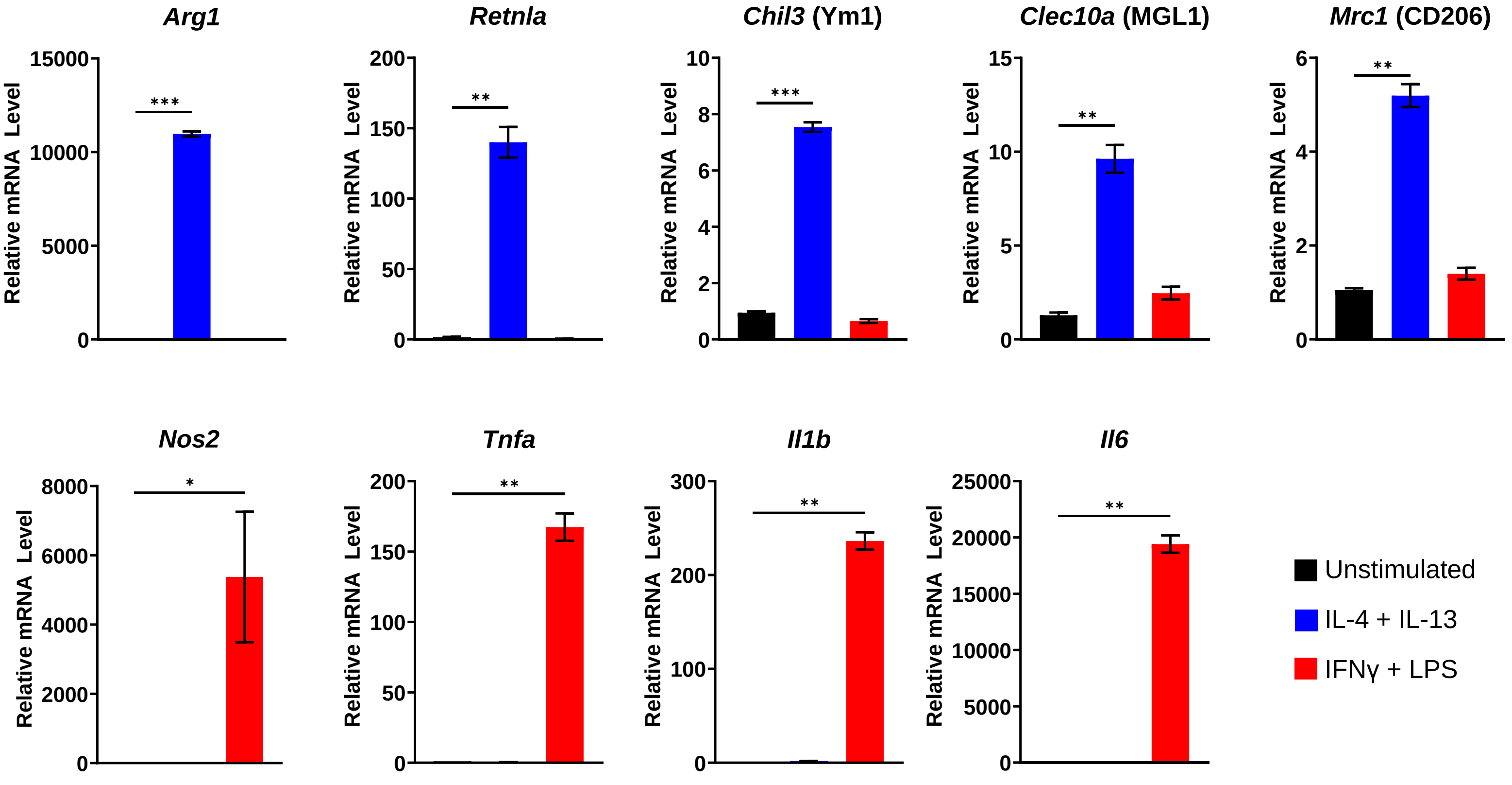

### Supplementary Fig 2

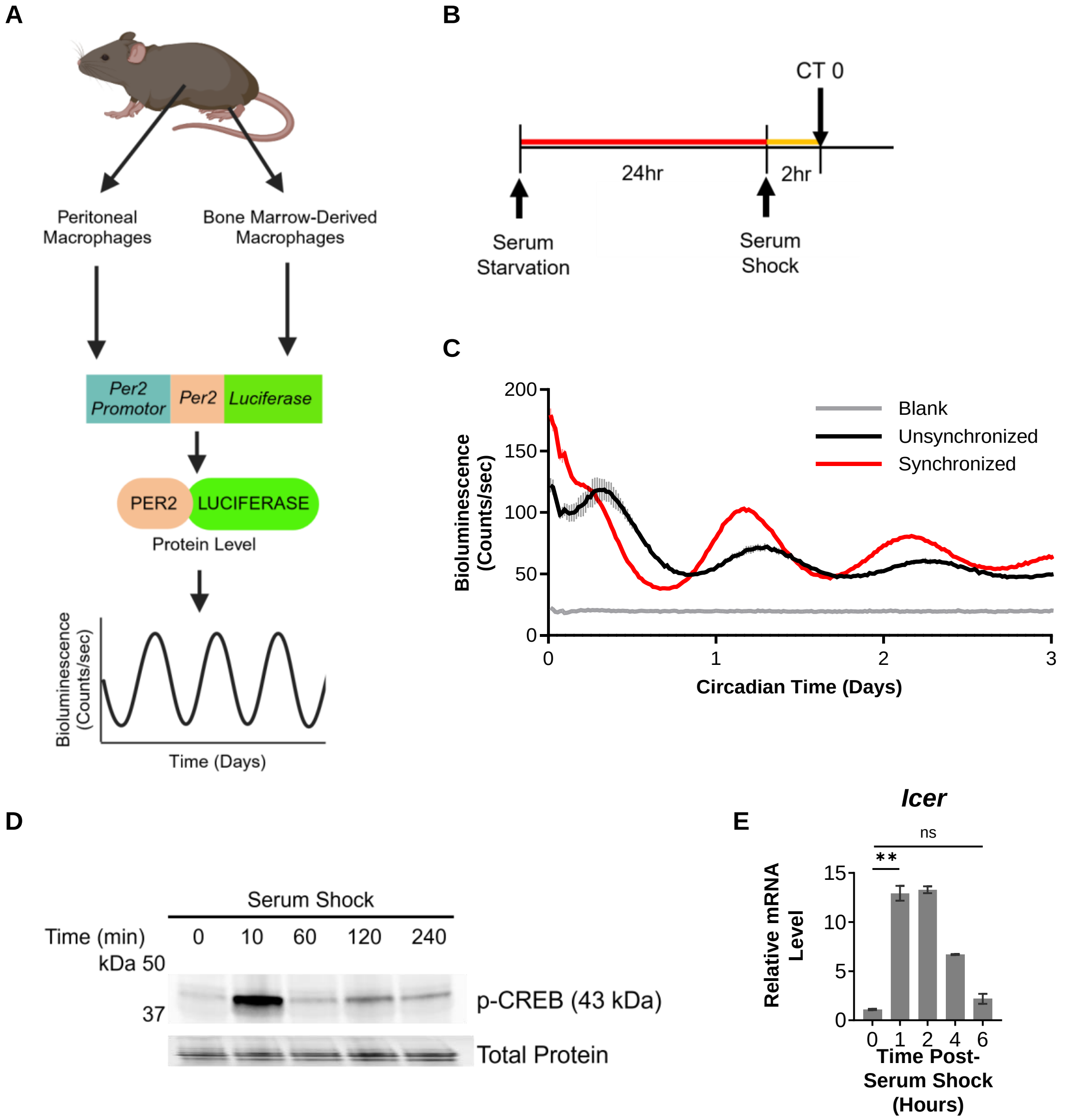

### Supplementary Fig 3

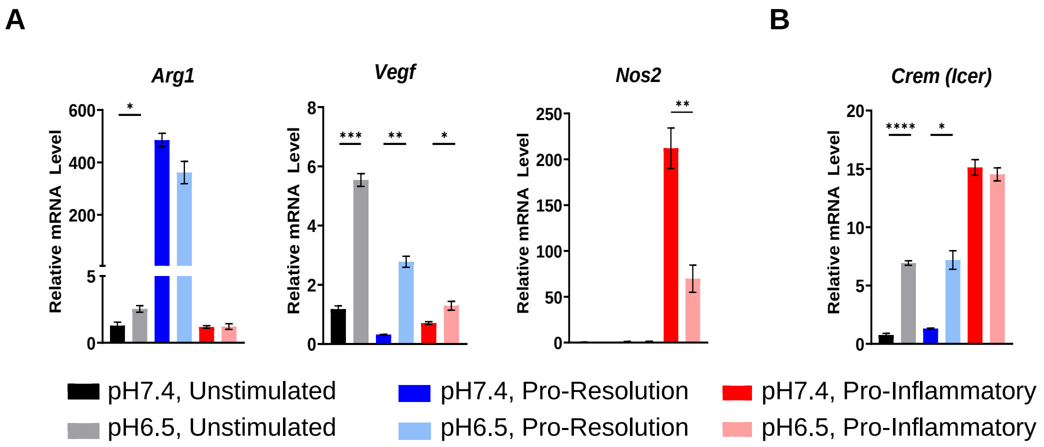

### Supplementary Fig 4

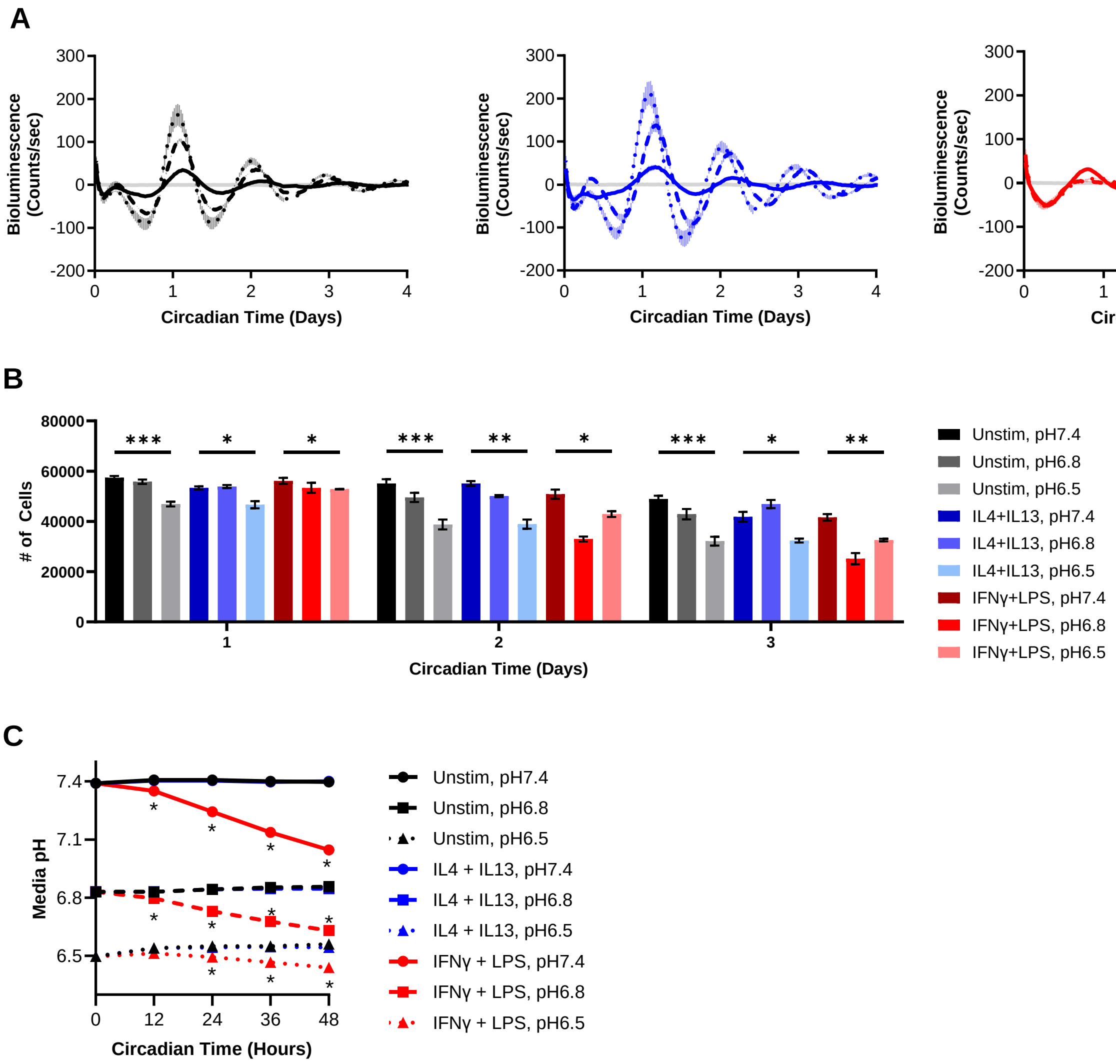

### Supplementary Fig 5

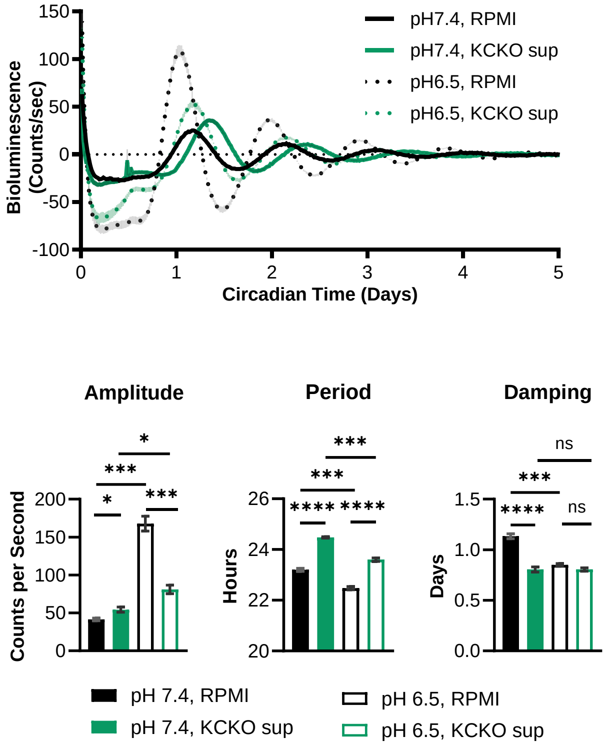

### Supplementary Fig 6

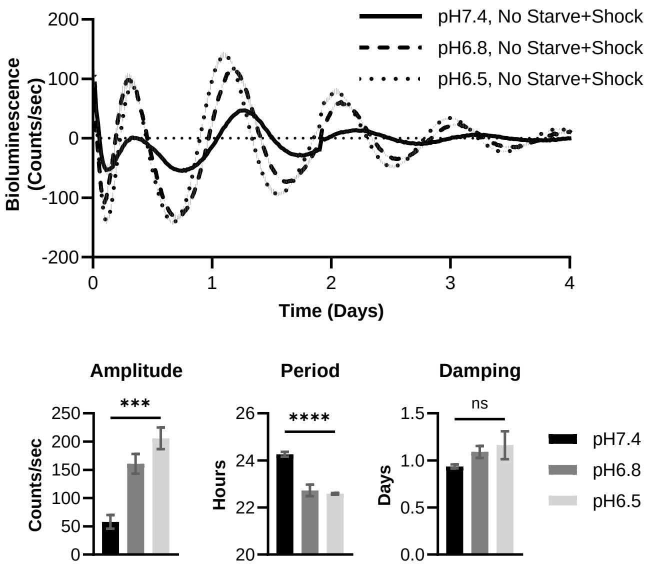

### Supplementary Fig 7

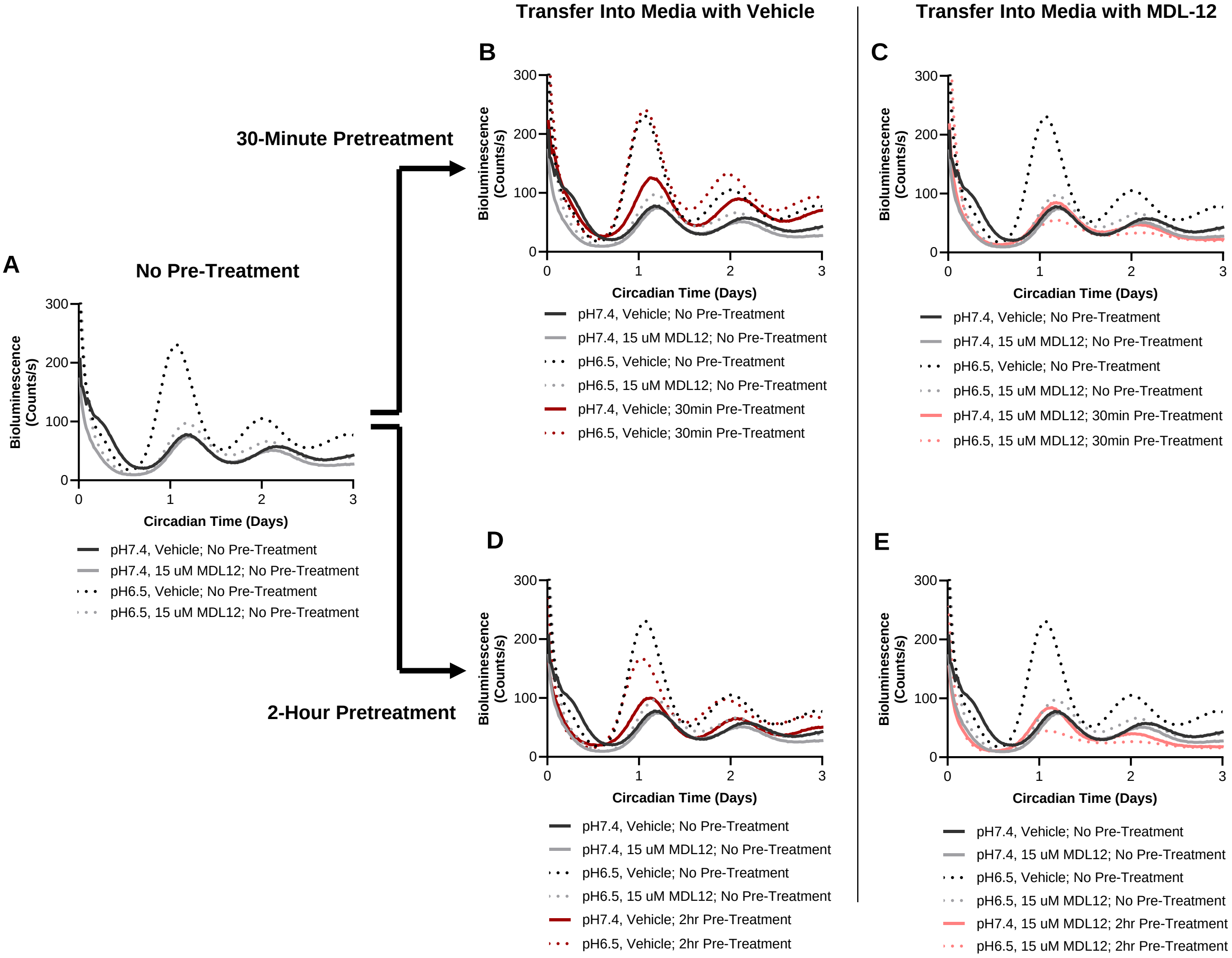

### Supplementary Fig 8

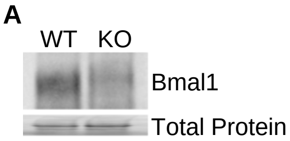


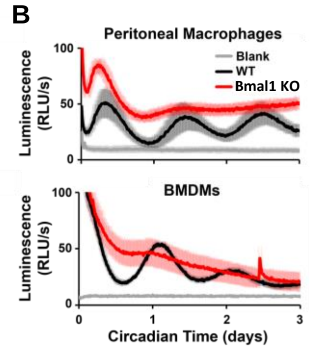

### Supplementary Fig 9

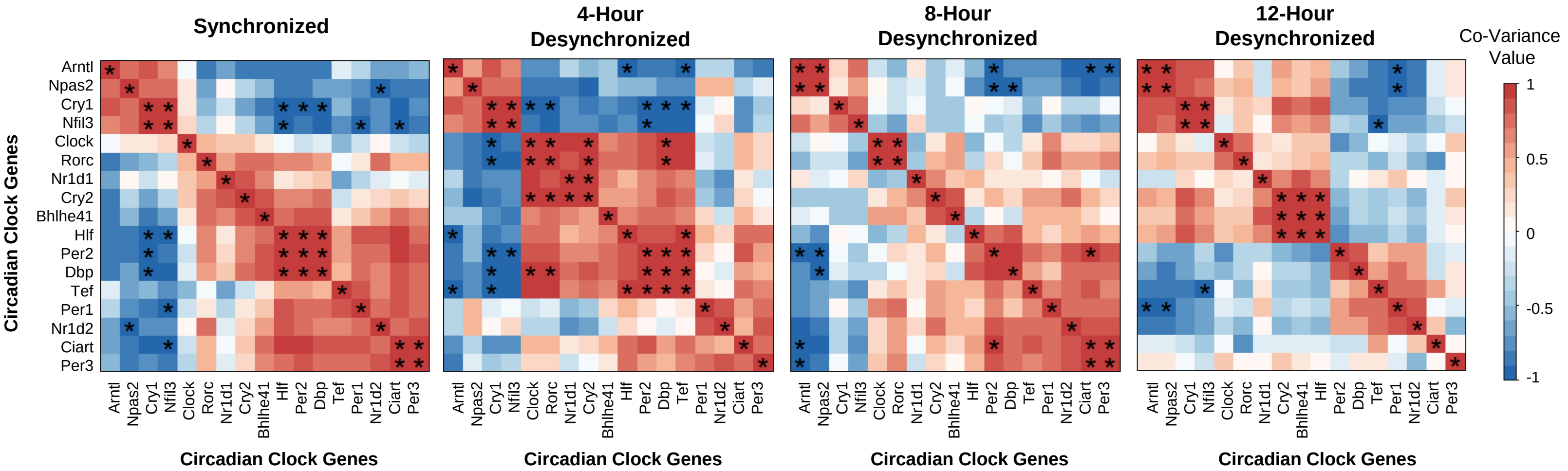
